# Exogenous attention interferes with endogenous attention processing via lateralized alpha power

**DOI:** 10.1101/2022.12.23.521791

**Authors:** Mathieu Landry, Jason Da Silva Castanheira, Amir Raz, Sylvain Baillet, Jérôme Sackur

## Abstract

Current theories of attention differentiate exogenous (bottom-up) from endogenous (top-down) orienting of visuospatial attention. While both forms of attentional processing engage different processes, endogenous and exogenous attention are thought to share processing resources, as shown by recent empirical evidence of their functional interactions. Here, we aim to uncover the neurobiological basis of how salient events that drive exogenous attention disrupts endogenous attention processes. We hypothesize that interference from exogenous attention over endogenous attention involves alpha-band activity, a neural marker of visuospatial attention. To test this hypothesis, we contrast the effects of endogenous attention across two experimental conditions: a single cueing condition where endogenous attention is engaged in isolation, and a double cueing condition where endogenous attention is concurrently engaged with exogenous attention. Our results are consistent with previous work and show that the concurrent engagement of exogenous attention interferes with endogenous attention processes. Importantly, we evaluate our main hypothesis using a moderated mediation model. We found that changes in alpha-band activity mediate the relationship between endogenous attention and its benefits on task performance, and that the interference of exogenous attention on endogenous attention occurs via the moderation of this indirect effect. Altogether, our results substantiate a model of attention, whereby top-down and bottom-up attentional processes compete for shared neurophysiological resources. This model accounts for the observed patterns of interference between exogenous and endogenous attention.

**Significance Statement:** Scientists differentiate top-down and bottom-up visuospatial attention processes. While bottom-up attention is rapidly engaged by emerging demands from the environment, top-down attention reflects slow voluntary shifts of attention. Several lines of research substantiate the idea that top-down and bottom-up attentional processes involve distinct brain systems. An increasing number of studies, however, argue that both attention systems share brain processing resources. The current study examines how salient visual events that engage bottom-up processes interfere with top-down processes. Using neurophysiological recordings and multivariate pattern classification techniques, the researchers show that both top-down and bottom-up attention processes do share brain resources expressed via alpha-band neurophysiological activity (8-12 Hz). The results further demonstrate that the interference patterns observed over brain activity in the alpha-band between both attention systems explains, in part, the interference between top-down and bottom-up attention at the behavioural level. The authors conclude by proposing a model of visuospatial attention whereby the dynamics between both attention systems are determined by their competition for limited brain processing resources.

## Introduction

We rely on visuospatial attention to accomplish everyday tasks. For example, when reading a journal article, we use goal-oriented attention to process the information written on the page. However, this process can be interrupted by salient stimuli like a notification on our phone. In this way, salient events can engage attention resources and thereby interfere with ongoing goal-oriented attention. The present work aims to explore these interference effects by investigating shared resources between top-down and bottom-up attention processes.

Indeed, researchers often differentiate at least two modes of control of visuospatial attention: Endogenous (i.e., top-down) attention that reflects the voluntary control of attention and exogenous (i.e., bottom-up) orienting which involves the involuntary shift of attention triggered by salient sensory events (Carrasco, 2011). Studies show that exogenous and endogenous attention are distinct processes that map onto separable brain systems (Chica, Bartolomeo, & Lupiáñez, 2013). Endogenous attention, for example, is slow to engage a target stimulus, whereas exogenous attention occurs rapidly. Likewise, exogenous attention is uniquely characterized by a significant delay before re-engaging to the same stimulus twice – a phenomenon called inhibition of return (Klein, 2000). Cognitive load manipulations that draw on top-down resources, on the other hand, selectively impair endogenous attention, but not exogenous attention (Jonides, 1981). The prevailing view is therefore of a dichotomous account of visuospatial attention.

Several studies, however, substantiated instances where exogenous and endogenous attention processes interfere with each other (Grubb, White, Heeger, & Carrasco, 2015; MacLean et al., 2009). The abrupt onset of a salient event (i.e., exogenous attention), for example, can disrupt endogenous attention processes–like a notification on your phone in the middle of reading this paragraph (Yantis & Jonides, 1990). In the same vein, one study reports interactions between exogenous and endogenous attention when task demands are high, e.g., when a visual target is difficult to discriminate. Behavioral responses of participants in this context are compatible with an interaction between both attention systems (Berger, Henik, & Rafal, 2005). Here, the occurrence of task-irrelevant stimuli can capture attention and cause slower response times. Evidence from electroencephalography (EEG) further corroborates these findings. During visual processing, both endogenous and exogenous attention alter the target-related N1 component, thereby revealing that they are not fully separable (Busse, Katzner, & Treue, 2008; Hopfinger & West, 2006; Landry, Silva Castanheira, Baillet, Sackur, & Raz, 2022). In sum, previous research suggests that both attention processes may share common neurophysiological resources from sensory systems, which may cause functional interferences.

In the present study, we hypothesize that interference of exogenous attention over endogenous attention would involve alpha-band (8-12 Hz) brain activity. This hypothesis follows from previous work showing that both attention systems express similar patterns of lateralized alpha-band brain activity (Foxe, Simpson, & Ahlfors, 1998; Keefe & Störmer, 2021; Störmer, Feng, Martinez, McDonald, & Hillyard, 2016). Alpha-band activity is thought to reflect modulations of cortical excitability, a central neural mechanism for the selection of attended sensory inputs (Klimesch, 2012). Numerous findings support this viewpoint, including evidence from a recent study showing that changes in alpha power in the occipital cortex plays a causal role in visuospatial attention (Bagherzadeh, Baldauf, Pantazis, & Desimone, 2020). In Bagherzadeh and colleagues’ study, participants were trained to alter lateralized patterns of alpha power over the posterior regions of the brain using neurofeedback. Critically, neurofeedback training was done independently from task-based shifts of visuospatial attention. Participants showed overall shifts of visuospatial attention thereafter. This implies that changes in alpha-band activity led to shifts of visuospatial attention, not the converse. Thus, we hypothesized that both exogenous and endogenous attention rely on this shared alpha-band neurophysiological signaling to amplify and process sensory inputs.

In line with this hypothesis, we previously reported evidence for the influence of endogenous attention on exogenous orienting over the power and phase of alpha-band activity (Landry et al., 2022). Here, classifiers exclusively trained to decode the effects of exogenous attention based on alpha rhythms were largely influenced by endogenous orienting when the same classifiers were tested in an experimental context comprising both attention processes. These findings corroborate the hypothesis that both attention systems share common neurophysiological processing resources (i.e., alpha-band activity). The present study aims to extend this research trajectory and investigate the influence of exogenous orienting on endogenous attention, i.e., investigate how salient events that capture attention via stimulus-driven processes can disrupt endogenous attention.

To test our hypothesis, we contrasted the effects of endogenous attention across two experimental conditions: a single-cueing condition where endogenous attention is engaged in isolation, and a double cueing condition where endogenous attention is concurrently engaged with exogenous attention. In this effort, we first confirmed that exogenous orienting interferes with endogenous attention over response times (i.e., behaviour) and target-locked ERPs (i.e., neurophysiology). Next, we tested whether the observed behavioural interference between endogenous and exogenous attention would be explained by alpha-band (8-12 Hz) EEG signal features expressed by both attention systems. We relied on a moderated mediation analysis to show that changes in alpha power mediate the relationship between endogenous cueing and task performance, while the exogenous cue moderates this mediation effect. Altogether, we propose a model of attention whereby the competition for shared neurophysiological resources leads to interference between exogenous and endogenous attention.

## Results

We collected data from thirty-two participants who completed three spatial cueing tasks in two experimental conditions, organized in a blocked design. Here, we present the results from two tasks: A single endogenous cueing task, which only consisted of an endogenous cue, and a double cueing task which consisted of both an endogenous cue and an exogenous cue. Note, results from our third task are presented in our previous report (Landry et al., 2022). In both conditions, participants were asked to discriminate the orientation of a Gabor target (clockwise versus counter-clockwise) presented at a cued or uncued location (Figure 1). We contrasted task performances for valid (i.e., target onset at the cued location) versus invalid (i.e., target onset at the uncued location) trials to estimate the benefits of endogenous and exogenous attention (i.e., cue validity effect, see Methods; Chica, Martín-Arévalo, Botta, & Lupiánez, 2014). Importantly, within the double cueing condition, we contrast trials where both the exogenous and the endogenous cues indicate the same location (i.e., congruent cueing trials), and where the cues indicate opposite locations (i.e., incongruent cueing trials). Hence, in the double cueing condition, we will use the term “congruent” to describe trials where both cues indicate the same location and “incongruent” to describe trials where the exogenous and endogenous cues indicate opposite locations (Figure 1).

**Figure 1.**
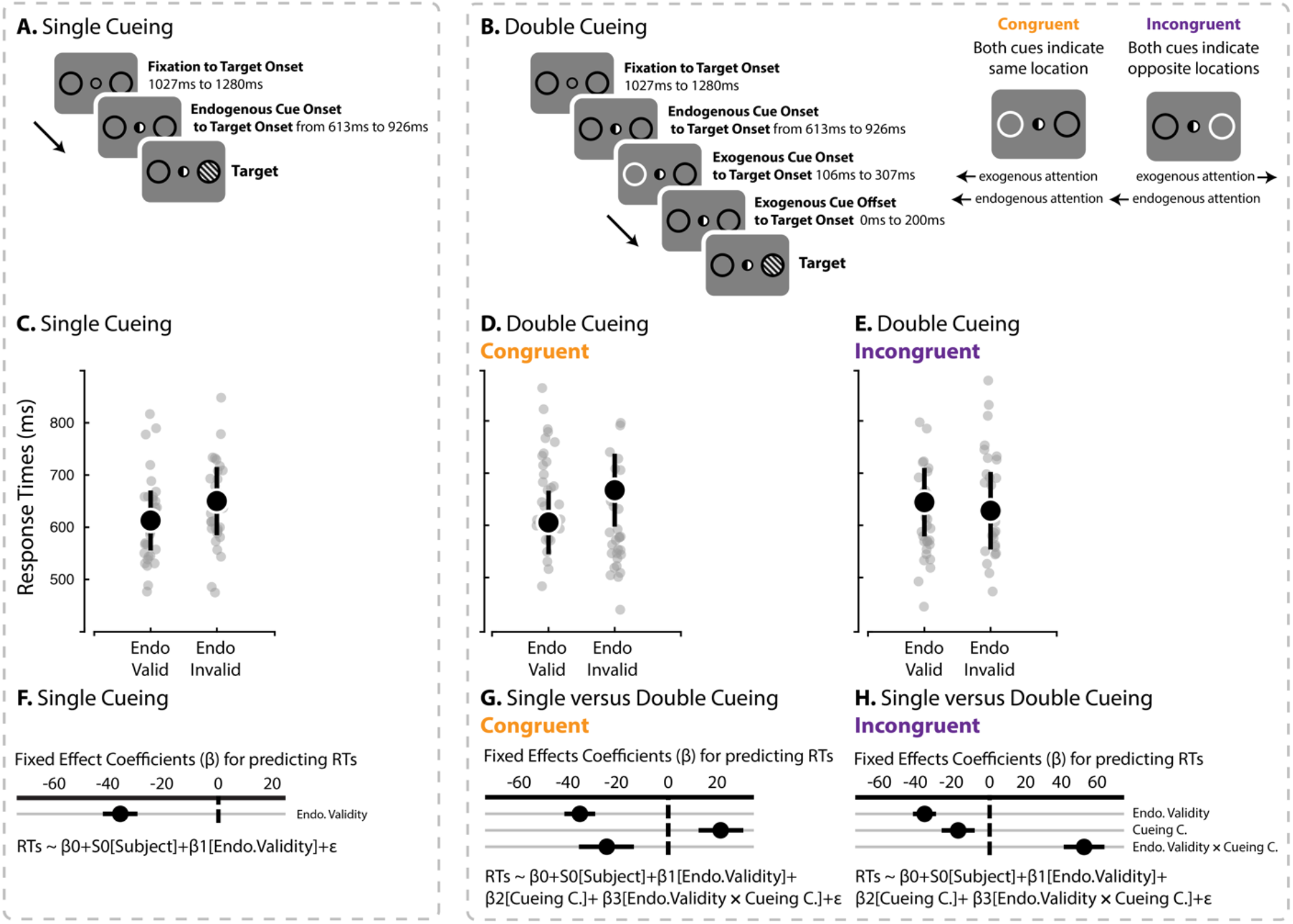
**A**. Single-cueing of endogenous attention. Each trial began with a fixation screen, followed by the onset of the endogenous cue after a random interval. The endogenous cue remained on screen until the end of trial. Next, a Gabor stimulus was shown at the left or right location. **B**. Double cueing condition comprising both exogenous and endogenous cueing. The trials began with a fixation screen followed by the endogenous cue, then the exogenous cue, and finally the target event onset at the left or right location. The endogenous cue was predictive of the target’s location in both cueing conditions, while exogenous cue validity was set to chance level. For both conditions (**A** and **B**, each trial ended with the participant’s discrimination response for the orientation of the Gabor stimulus (clockwise versus counter-clockwise). In both cueing conditions, onsets of the endogenous cue, the exogenous cue, and the target stimulus were time jittered. Diagram at the top right corner shows that the double cueing condition divides into trials where both cues indicated the same location (i.e., congruent cueing condition) and trials where both cues indicated opposite locations (i.e., incongruent cueing condition). The plots show groups (black dots) and participants (grey dots) averaged accurate RTs for endogenous cue validity across single cueing (**C**), double cueing congruent trials (**D**), and double cueing incongruent trials (**E**). Error bars represent bootstrapped 95% C.I. Bottom graphs indicate single-trial hierarchical regression model coefficients and corresponding 95% C.I. For single cueing (**F**), we included endogenous cue validity (dummy coding was 0 for cue invalid and 1 for cue valid) as a fixed predictor for predicting correct RTs. For double cueing, we evaluated endogenous cue validity across single versus double cueing for congruent cueing condition (**G**) and incongruent cueing condition **(H)**. In both instances, we included endogenous cue validity, cueing condition (i.e., single versus double) and their interaction as fixed factors.

### Behavior

The participants identified the target orientation with near-ceiling performances (93% average accuracy rate across participants). Consequently, our behavioral analyses focused on accurate response times (RT). Our behavioral analysis was twofold. First, we wanted to confirm that the endogenous and exogenous cues influenced performance in the single cueing (i.e., endogenous cue alone) and double cueing (i.e., endogenous and exogenous) conditions. We then tested if exogenous attention impacted the benefits of endogenous attention by contrasting the endogenous cue validity effect across single and double cueing conditions (i.e., when the exogenous cue is present versus when it is absent).

We confirmed that the exogenous and endogenous cues facilitated performance in both cueing conditions. In the single-cueing condition, endogenous attention decreased mean RT by 36 ms (endogenous cue validity effect; β=−36.49, SE=3.24, 95% CI [−42.84, −30.13]; Supplementary Tables 1 and 2). Likewise, in the double cueing condition, we observed main effects of exogenous and endogenous cue validity whereby participants were 39 ms faster for valid exogenous cues (exogenous cue validity effect; β=−39.27, SE=3.04, 95% CI [−45.22, −33.32]) and 22 ms faster for valid endogenous cueing (endogenous cue validity effect; β=−22.42, SE=3.22, 95% CI [−28.73, −16.11]; Supplementary Table 4). Evidence weighted against the interaction between endogenous cue validity and exogenous cue validity in the double cueing condition (Bayes factor, BF_01_=105.59). Thus, both cues produced their expected effects.

Next, we confirmed that the presence of the exogenous cue altered the endogenous cueing effect by contrasting the cue validity effects of endogenous attention in the single cueing vs. double cueing conditions. We performed this analysis across congruent and incongruent cueing conditions (Figure 1). For congruent trials, we observed a significant interaction between endogenous cue validity and cueing condition (β=−25.2, SE = 5.68, 95% CI [−36.33, −14.06]; Figure 1C, and Supplementary Tables 5 and 6) that indicated that the cueing effect of endogenous attention increased from single and double cueing. For these trials, exogenous attention boosted endogenous attention. Conversely, for incongruent trials, our results indicated that exogenous attention interfered with the cueing effect of endogenous attention, as evidenced by a significant interaction between endogenous cue validity and cueing condition (β = 52.74, SE = 5.72, 95% CI [41.52 63.97]; Figure 1D, and Supplementary Tables 7 and 8). These results provide evidence of interference from exogenous attention over endogenous attention whenever they are engaged towards opposite locations.

### Electrophysiology: Exogenous attention disrupts the cueing benefits of endogenous attention over target-locked event-related potentials

Next, we aimed to corroborate the interference effect of exogenous attention at the neural level using a multivariate pattern classification approach adapted from Bae and Luck (2018). Previous work highlights the benefits of this approach for studying visuospatial attention (Hong, Bo, Meyyappan, Tong, & Ding, 2020; Landry et al., 2022). Consistent with our behavioral results, we expected that exogenous attention would interfere with target-related processing.

We first trained linear support vector machine (SVM) classifiers at each timepoint of the target-locked event-related potentials (ERP) using a three-fold cross-validation procedure in the context of single cueing. The SVM classifiers were trained to separate endogenous cue valid ERP from endogenous cue invalid ERP across the time series. We determined the classification performance by evaluating the classifiers’ capacity to accurately classify ERP on the left-out ERP time-series (i.e., the validation set). We therefore trained and tested our SVM linear models in the context of single cueing. To examine the interference of exogenous attention, we then tested these SVM models in the context of double cueing for congruent and incongruent trials separately. See Figure 2 and Method section for details.

**Figure 2.**
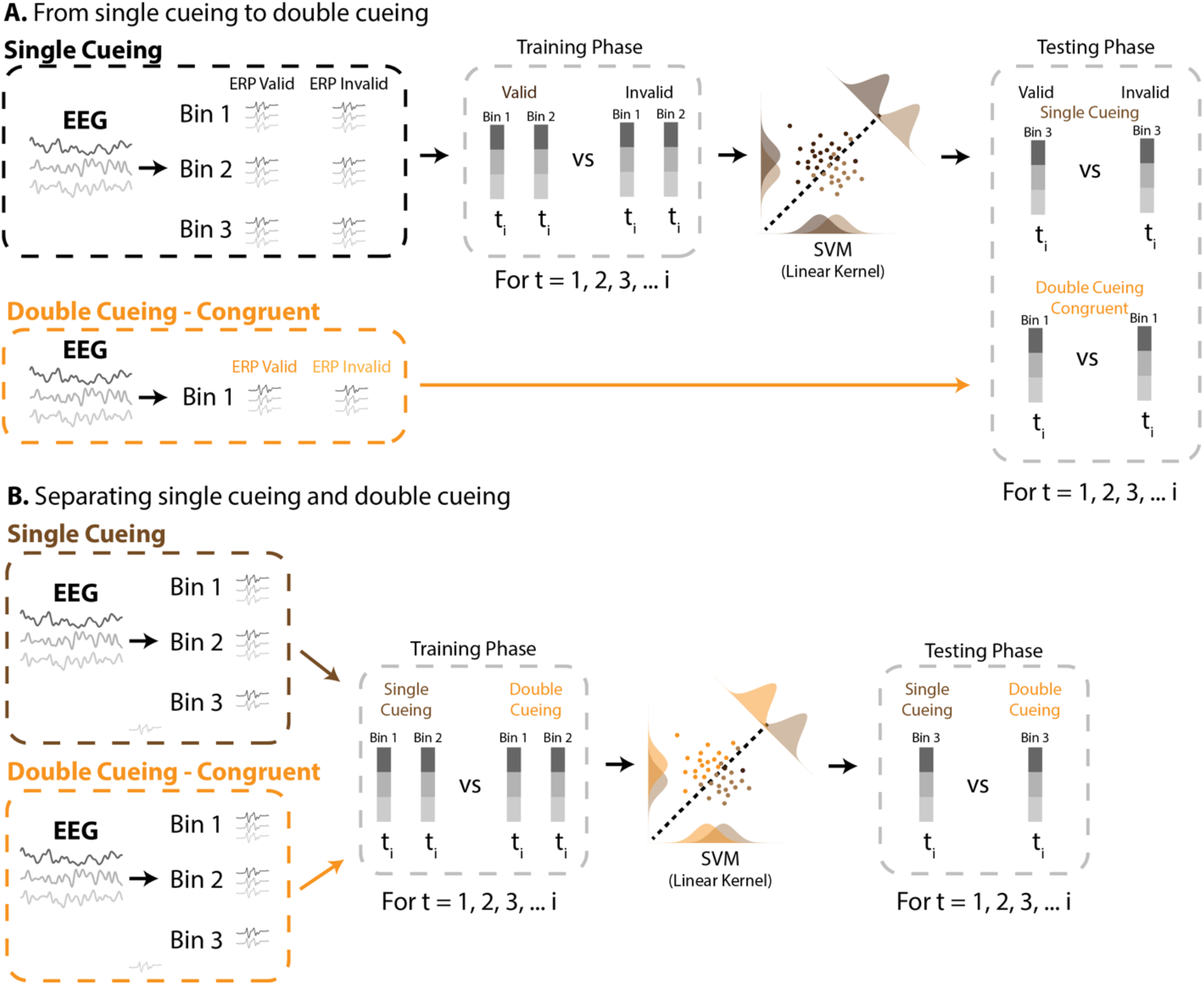
**A**. Linear SVM classification procedure to decode endogenous cue validity. We used a three-fold cross-validation procedure whereby we trained classifiers at each time point using two ERP for valid trials and two for invalid ones from the single cueing condition, and then tested the same classifiers on a third ERP for valid trials and invalid trials from the single cueing condition. Next, we assessed the performance of these SVM classifiers in the context of double cueing. In this example, we exhibit how classifiers trained during single cueing are tested in the context of double cueing congruent trials. The same procedure was used when we tested these models during double cueing incongruent trials. **B**. Linear SVM classification to decode single versus double cueing. Again, we used a three-fold cross-validation procedure whereby we trained classifiers at each time point of two EEG time series from single- and double cueing conditions, and then tested these models using left-out data from a third time series. In this example, we exhibit how classifiers trained and tested to decode single cueing vs. double cueing congruent trials. The same procedure was used when we evaluated single cueing vs. double cueing incongruent trials.

We first validated the performance of classifiers trained in the single-cueing condition to decode endogenous cue validity (i.e., valid versus invalid trials) based on target-locked event-related potentials (ERPs). The SVM classifiers performed above-chance level between 112 and 388ms following target onset (Figure 3A), which confirmed the ability of the models to separate valid and invalid trials during this period. We then evaluated how well these classifiers would perform when tested in the context of double cueing. We reasoned that poorer classification performances in the double cueing condition would indicate the effect of an interference of exogenous attention over endogenous attentional processes. Decoding was successful for congruent cueing trials, as evidenced by several significant clusters starting at 160ms post-target onset and that lasted almost the entire epoch (Figure 3B). Conversely, for incongruent cueing trials, the decoding of endogenous cue validity was unsuccessful while the classifiers performed at chance-level, and even below it between 256 and 328ms (Figure 3C). These results are consistent with our behavioral findings and show that when a salient event occurs to the opposite location relative to endogenous attention prevents the classification of endogenous cue validity from target-locked ERP. The comparison of decoding performance across the 3 conditions (i.e., single cueing, double cueing congruent trials, double cueing incongruent trials): We did not observe any difference between single cueing and double cueing congruent trials, whereas we found significant clusters when we compared single cueing between double cueing incongruent trials (see supplementary Figure 1).

**Figure 3.**
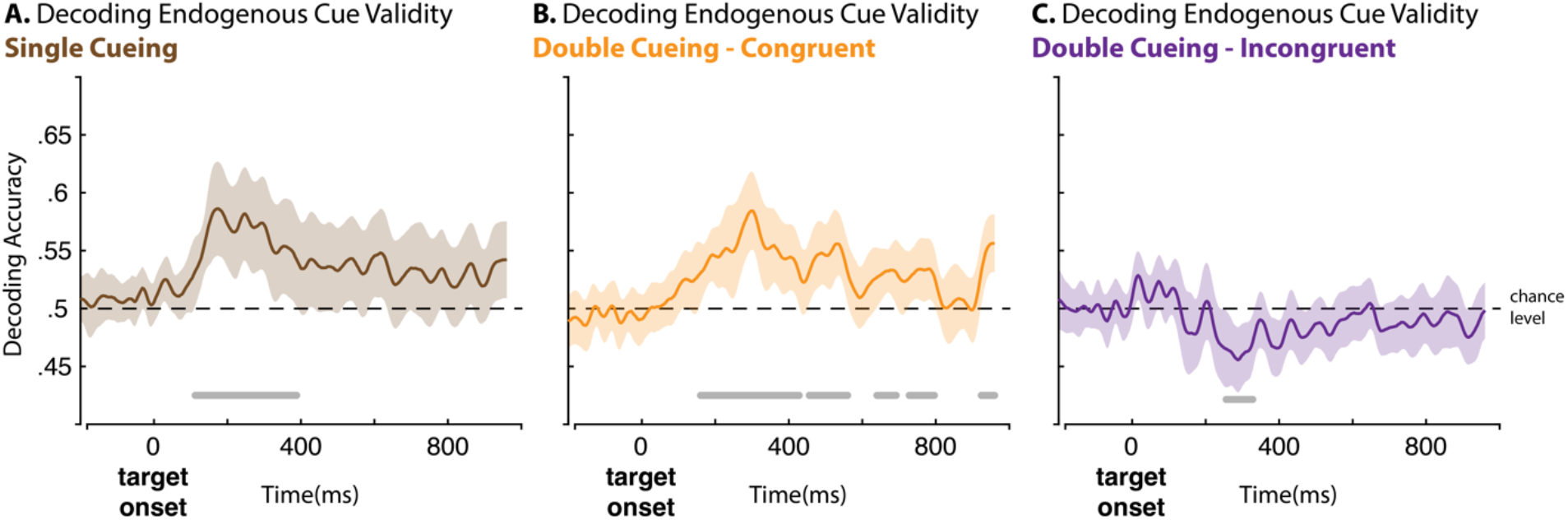
Group averages of decoding accuracy from target-locked ERP in the single-cueing **(A)**, double cueing congruent cueing trials **(B)**, and double cueing incongruent cueing trials **(C)**. SVM classifiers were trained across for each time point of the time series to separate endogenous cue valid trials from endogenous cue invalid trials during single cueing. We then tested this model on left-out data from the single cueing condition (**A**), on double cueing congruent cueing condition (**B**), and double cueing incongruent cueing condition (**C**). We used one sample t-tests, random cluster permutation and mass t-statistics to test for significant differences in classification performances against chance-level (50%). Horizontal grey lines indicate the temporal segments of significant differences in decoding accuracy. Shaded areas represent the 95% C.I.

We examined target-locked ERPs to better understand how the onset of exogenous attention to the opposite location impacts endogenous cue validity. Our approach follows from previous work that highlights the influence of visuospatial attention on the P1-N1 complex at the contralateral side (Hopfinger & West, 2006; Luck, Woodman, & Vogel, 2000). We averaged the waveforms at channels PO7 and PO8 based on target-location across single cueing endogenous cue valid trials, single cueing endogenous cue invalid trials, double cueing incongruent endogenous cue valid trials and double cueing incongruent endogenous cue invalid trials. This approach yielded four separate ERPs. We then compared these waveforms across the time series using hierarchical regressions at each time point where we included cue validity (endogenous cue valid vs. endogenous cue invalid) and cueing condition (single- vs. double cueing) and their interaction as fixed factors and participants as random factors. We cluster-corrected based on cluster-mass. This analysis returned a significant cluster for the interaction between the two variables at the time of the P1-N1 complex, thereby confirming that the onset of the exogenous cue altered the effect of endogenous attention over early visual processing (Figure 4).

**Figure 4.**
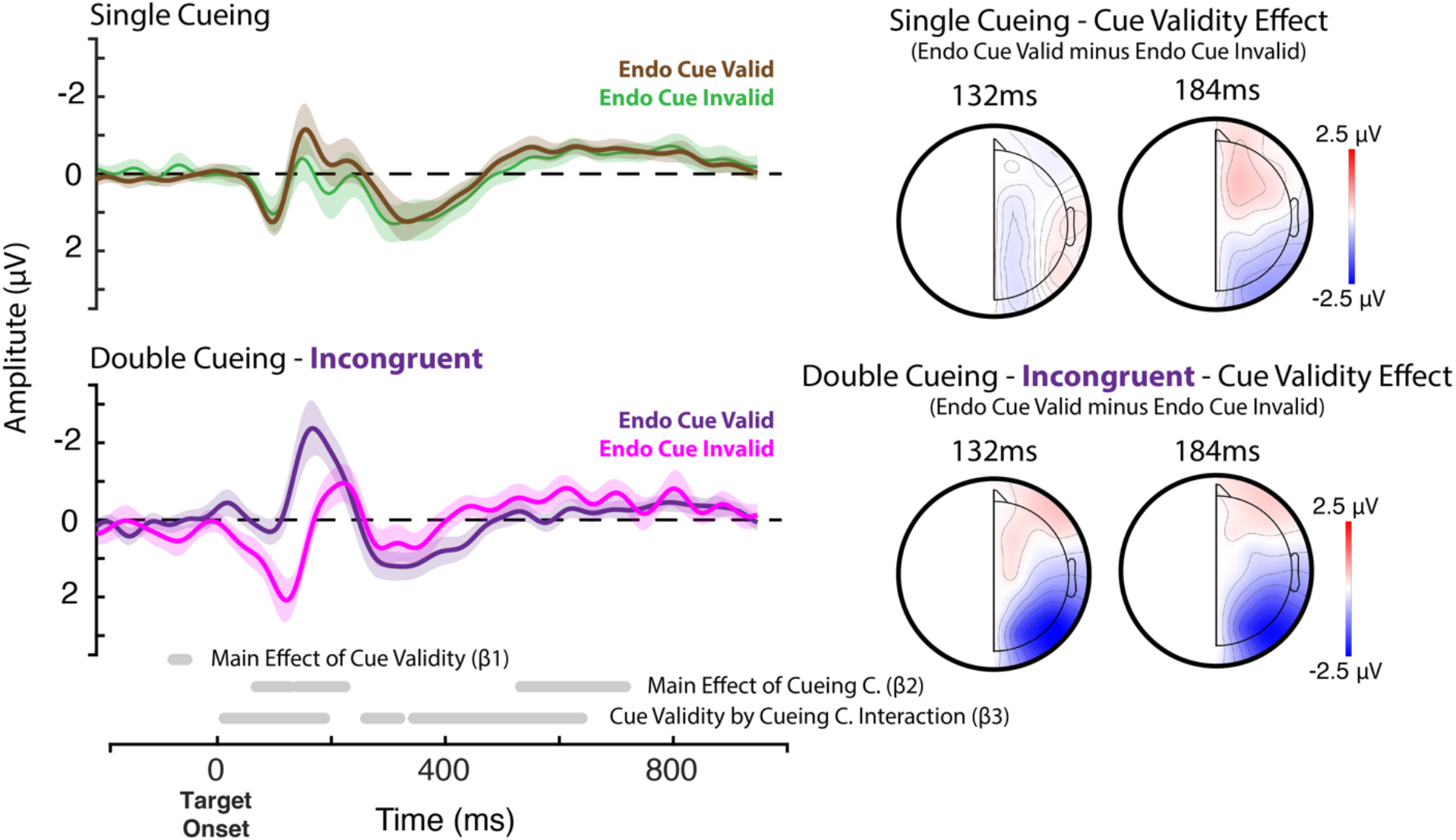
Target-locked averaged waveforms for endogenous cue valid and endogenous cue invalid trials at the contralateral side relative to the location of the target for channels PO7 and PO8. Line represents group averaged waveforms, shaded area represents the 95% C.I. We extracted waveform from channel PO8 when the target even occurred on the left side of the screen, and channel PO7 when it occurred on the right side. Top-row shows the data from the single cueing condition, bottom row data from the double cueing incongruent. We examined whether the endogenous cue validity effect varied as a function of cueing condition using the following hierarchical regression model across the time series: Amplitude ~ 1 + (1|participants) + β1[Endogenous Cue Validity] + β2[Cueing Condition] + β3[Endogenous Cue Validity × Cueing Condition] + ε. We cluster-corrected the regression coefficients based on mass t-statistics. Significant clusters across the time series are shown at the bottom. Right side of the figure shows the corresponding topographies for the endogenous cue validity effect: endogenous cue valid minus endogenous cue invalid for single cueing (top) and double cueing – incongruent (bottom).

Furthermore, we aimed to confirm that the mere presence of the exogenous cue to the opposite location of endogenous attention (i.e., incongruent cueing trials) interferes with the engagement of endogenous attention processes toward a target event. To evaluate this question, we solely focused on endogenous cue valid trials (i.e., those trials where the target event happened at the endogenous cued location). Here, we tested the capacity of our SVM classifiers to accurately separate endogenous cue valid trials of the single cueing condition from trials of the double cueing - incongruent cueing condition (Figure 2B). We adopted the same three-fold cross-validation approach from our previous decoding analysis and used target-locked ERP. We investigated ipsi- and contralateral channels relative to the location of the target separately. Our results confirmed that the onset of the exogenous cue to the opposite location alters the processing of a target event by endogenous attention. For contralateral channels, we observed two significant clusters: A first one from 96 to 604ms, and then another from 628 to 744ms (Figure 5A). A very similar pattern emerged for ipsilateral channels where we observed three significant clusters: A first one from 80 to 620ms, another from 656 to 728ms, and from 782 to 860ms (Figure 5B). In short, the presence of the exogenous cue to the opposite location altered the processing of the target event by endogenous attention throughout almost the entire time series.

**Figure 5.**
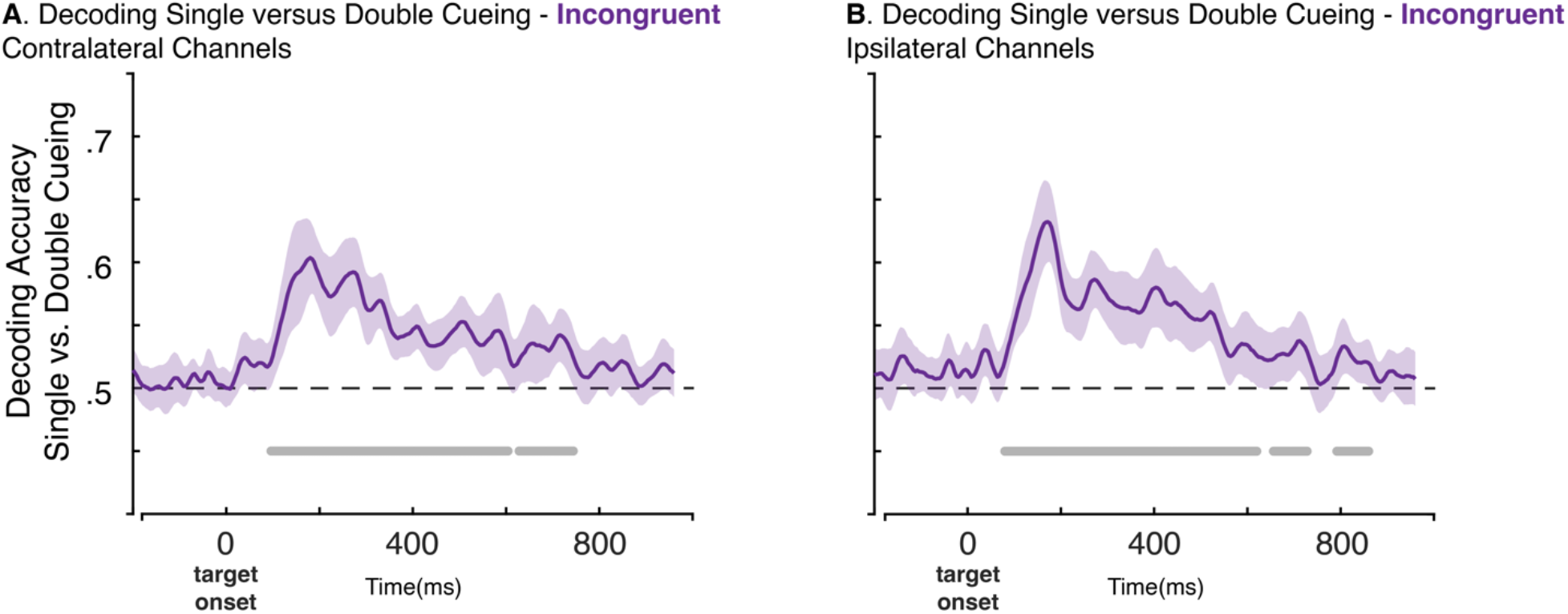
Group averages of decoding accuracy from target-locked ERP for endogenous cue valid trials only. We performed separate analysis for ipsilateral (**A**) and contralateral (**B**) channels relative to target location. SVM classifiers were trained across each time point of the time series to separate single cueing from double cueing incongruent cueing trials. We tested these classifiers on left-out data using one sample t-tests, random cluster permutation and mass t-statistics to test for significant differences in classification performances against chance-level (50%). Horizontal grey lines indicate the temporal segments of significant differences in decoding accuracy. Shaded areas represent the 95% C.I.

### Electrophysiology: Modulations of alpha power distinguishes between single & double cueing

We previously described how alpha-band activity plays a central role in visuospatial attention and hypothesized that interference of exogenous attention over endogenous attention would involve this neural feature. To test our hypothesis, we first trained classifiers to separate cueing conditions (i.e., single versus double cueing; Figure 2B) from the lateralization of alpha-band brain activity (LAI) relative to the orienting of endogenous attention. We therefore aimed to confirm that alpha-band activity varied between single and double cueing conditions. For this analysis, we target-locked the time series and focused on the period that precedes the target-stimulus. We did not observe any significant cluster around exogenous cue onset for decoding the cueing condition when cues pointed to the same location in the double cueing condition (Figure 6A). Conversely, for incongruent cueing trials of the double cueing condition, classification performance of cueing condition was above chance-level from −336ms to 192ms relative to target onset (Figure 6B). This outcome supports our hypothesis and shows that the presence of the exogenous cue to the opposite location of endogenous orienting modulates alpha-band activity pertaining to endogenous attention. Importantly, the corresponding decoding weights at that time highlighted posterior contributions (Figure 6D). Unexpectedly, we also found a significant cluster around the onset of the endogenous cue (Figures 6B), which implies that the endogenous cue was processed differently in the single and double cueing conditions, most likely reflecting a contextual effect related to our block design. We, however, observed a significant cluster around the onset of the endogenous cue.

**Figure 6.**
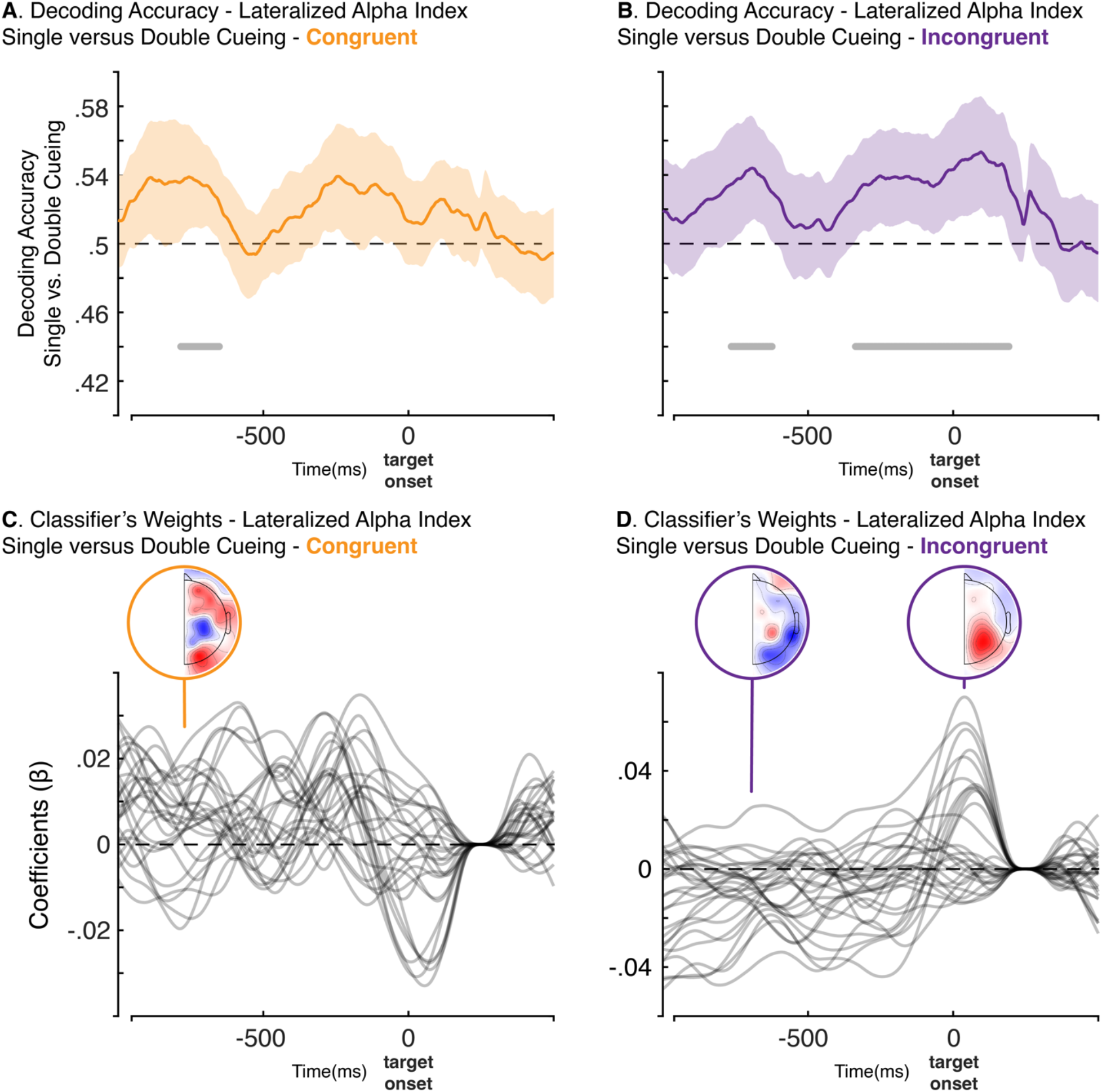
Group averages of decoding accuracy from target-locked changes in lateralized alpha power (LAI) relative to the orienting direction of endogenous attention. SVM classifiers were trained on each time point of the time series to separate single-cueing from double cueing congruent trials (**A**) and single-cueing from double cueing incongruent trials (**B**). We tested these models on left-out data using one sample t-tests, random cluster permutation and mass t-statistics to test for significant differences in classification performances against chance-level (50%). Horizontal grey lines indicate the temporal segments of significant differences in decoding accuracy. Shaded areas represent the 95% C.I. Bottom-up plots show the corresponding averaged SVM coefficients across the time series, as well as their topographies (**C, D**).

### Interference in alpha-band power explains behavioural interference between attention systems

Our results highlight the interference of exogenous attention over endogenous attention whenever both cues indicate opposite locations, both at the behavioral and neural levels. We also related the effects of the exogenous cue over endogenous attention to the modulations of lateralized alpha power. Our final analysis aimed to link and examine whether the interference effect of exogenous attention over endogenous attention that we observed at the behavioral level relates to changes in lateralized alpha power using a moderated mediation analysis. For this analysis, we solely focused on incongruent cueing trials. Here, we first extracted the target-locked LAI values at the time and for the channel where we had observed the largest SVM beta coefficients for decoding the cueing condition (i.e., single vs. double cueing incongruent). This means that we choose the time and channels where we observed the maximal effect of the exogenous cue on LAI, which corresponded to channels P3/P4 at 40ms post-target onset. Our moderated mediation model entails two hypotheses: A first one where changes in the LAI values we extracted would mediate the relationship between endogenous cue validity and RT, and then a second one where the onset of the exogenous cue for incongruent cueing trials would moderate this meditation.

We first verified that the effect of endogenous cue validity on RTs is mediated by changes in LAI. Here, we found evidence for a partial mediation effect (indirect effect: β=11.93, 95% CI [3.19, 20.67]), given that the direct effect remained statistically significant: (direct effect: β=−26.70, 95% CI [−44.4, −9.01]). See supplementary Figure 2 and supplementary tables 9 and 10. Next, we added the cueing condition (i.e., single vs. double cueing) as a moderator of this indirect mediation effect. The moderated mediation model confirmed our hypothesis and showed that incongruent cueing interferes with endogenous cue validity by moderating the mediation of lateralized alpha-band activity, (moderated mediation index: β=20.06, 95% CI [1.45, 43.66]; Figure 7). Furthermore, the moderation mediation effect of double cueing on the mediation path fully accounted for the interference of exogenous attention effect on behavior, whereby the interaction between endogenous cue validity and cueing condition for predicting RTs was non-significant in our moderated mediation model, β=18.54, 95% CI [−27.05, 55.28] (Figure 7). See supplementary tables 11 and 12.

**Figure 7.**
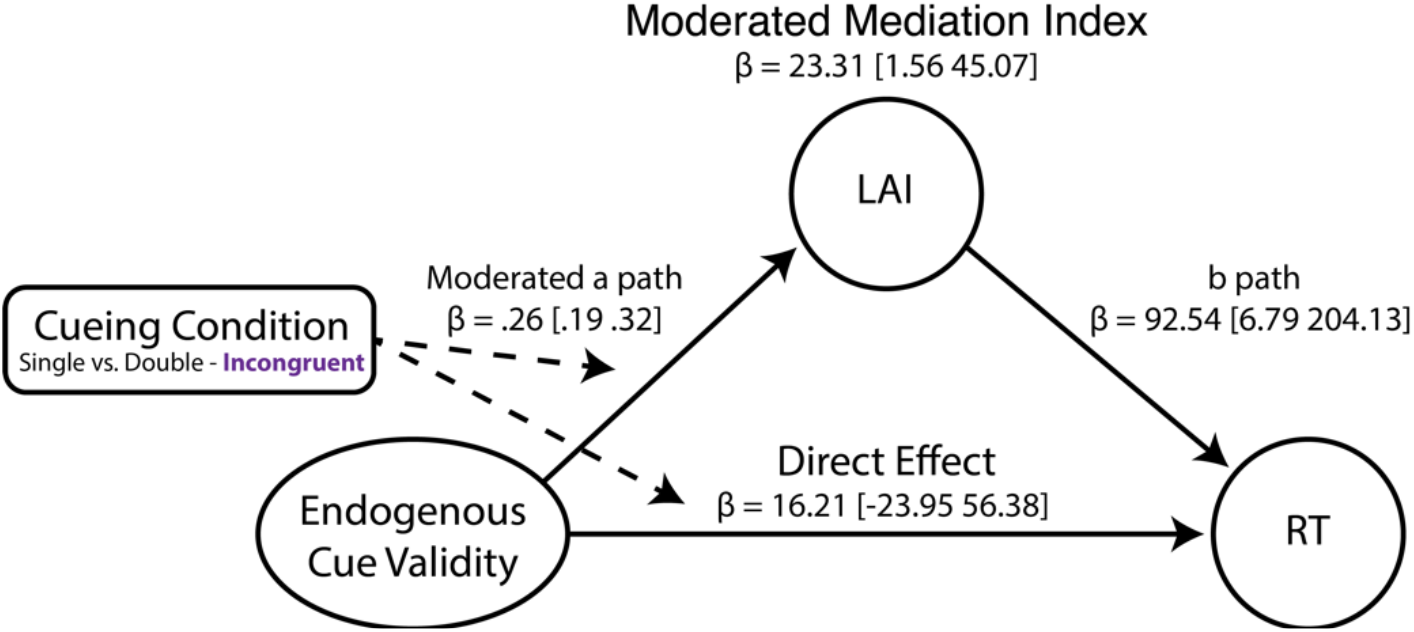
Moderated mediation analysis where the direct effect involves endogenous cue validity predicted response times (RT), while the cueing condition moderates this relationship. The indirect effect involves mediation through lateralized alpha power (LAI) at channels P3/P4 at 40ms post-target onset, wherein the cueing condition moderates the relationship between endogenous cue validity and LAI. The moderation effect of the cueing condition over the relationship between endogenous cue validity and response times is fully accounted for by the moderated mediation.

## Discussion

In the present work, we investigated the neurophysiological basis for the interference of exogenous attention on endogenous attention. To this end, we contrasted the effects of endogenous attention across two conditions: Single cueing where endogenous attention is engaged alone, and double cueing where endogenous attention is concurrently engaged with exogenous attention. This simple experimental approach confirmed that a salient peripheral cue that engages exogenous attention interferes with endogenous attention whenever both attention processes are directed to opposite locations. These findings are consistent with previous work showing that task-irrelevant salient events capture attention in a manner that can impede endogenous attention processes (e.g., Theeuwes, 1991). We observed this interference pattern both at the behavioral and neural levels. We hypothesized that these effects would relate to lateralized alpha-band activity – a neural marker of visuospatial attention (Keefe & Störmer, 2021; Störmer et al., 2016). Our mediated moderation analysis supports this hypothesis. We propose a model of visuospatial attention where the dynamics of exogenous and endogenous attention are governed by the competition for limited neural processing capacity (e.g., alpha rhythms).

Our findings confirmed that the concurrent engagement of exogenous attention influences endogenous attention (Figure 1). We observed this effect at the behavioral level where exogenous attention boosted the cueing effect of endogenous attention when both cues indicated the same location (i.e., congruent cueing trials), and then lessened the endogenous attention cueing effect when the cues pointed to opposite locations (i.e., incongruent cueing trials). While both attention processes correspond to distinct functional systems, this outcome validates that these systems do not fully operate independently from each other (Berger et al., 2005). We further corroborated these findings at the neural level by showing that cueing them to opposite locations significantly impacts the decoding of the endogenous cue validity based on target-locked ERPs, which dropped to chance-level (Figure 3C). Likewise, our study reveals that the engagement of endogenous attention towards target stimuli was also impaired independently from cue validity effects (Figure 5). These outcomes align with the idea of capacity-limited attention processing, whereby the advent of a salient event (i.e., the exogenous cue) prevents endogenous attention from fully engaging visual events (Alvarez & Franconeri, 2007; Busse et al., 2008). Our results are also consistent with past reports showing that interactions between exogenous and endogenous attention occur relatively early during sensory processing (Hopfinger & West, 2006; Landry et al., 2022). Here, we found that the presence of the exogenous cue to the opposite location impacted the P1-N1 complex at the contralateral site – two early ERPs modulated by visuospatial attention (Figure 4; e.g., Hopfinger & West, 2006). This early interaction pattern between endogenous and exogenous attention processes is compatible with neuroimaging findings relating both exogenous and endogenous attention to the primary visual cortex (Dugué, Merriam, Heeger, & Carrasco, 2020; Müller & Ebeling, 2008). The present work therefore demonstrates overlap between these attention processes over early visual processes.

Our main hypothesis derives from a large body work showing that shifts of visuospatial attention coincide with decreased amplitude of alpha power over the contralateral posterior region of the brain and increased amplitude for the ipsilateral region (Klimesch, 2012; Peylo, Hilla, & Sauseng, 2021). Prevailing views postulate that this lateralized pattern of alpha-band activity reflects changes in neural excitability and inhibition during visuospatial attention (Klimesch, 2012). Specifically, decreased alpha-band activity is thought to mark increased neural processing for events occurring in the attended visual field, whereas increased alpha power indexes the suppression of unattended sensory inputs. Consistent with this dominant viewpoint, we observed that the onset of the exogenous to the opposite location altered alpha rhythms related to endogenous orienting of attention (Figure 6B) – an effect that was related to the posterior region of the brain (Figure 6D). Conversely, we did not observe such changes in alpha-band activity when the exogenous cue occurred at the same location (Figure 6A). The onset of exogenous attention therefore altered lateralized alpha power activity related to endogenous attention, which corroborates that both attention processes involve modulations of this neurophysiological marker of visuospatial attention. In a previous study, we demonstrated that exogenous attention yielded significant changes of the power and phase of alpha rhythms (Landry et al., 2022). The current results therefore complement these previous findings. We then verified whether these effects related to alpha power relate to the interference effects we observed at the behavioral level. We used a moderated mediation analysis to test this main hypothesis relating the interference effect of exogenous attention over endogenous cue validity effect to lateralized alpha-band activity. Here, we found that cueing condition moderated the mediation path relating endogenous cue validity and response times via alpha-band activity (Figure 7). Specifically, the onset of the exogenous cue to the opposite location alters endogenous attention related effects on alpha-band activity. Importantly, this moderated mediation effect fully accounted for the interference effect of exogenous attention over endogenous cue validity we observed at the behavioral level. Our results therefore show that modulations of alpha-band activity are not only related to shift of visuospatial attention, but also relates to the perceptual benefits following attention processing. These findings support our main hypothesis and show that modulations of alpha-band synchrony reflect core mechanisms of visuospatial attention and that these mechanisms are shared between exogenous and endogenous processes, which leads to interference between them.

The present findings complement our recent work showing that endogenous attention overlaps with exogenous attention across alpha-band activity (Landry et al., 2022), and is consistent with our view that alpha rhythms reflect shared neural resources and capacity-limited processing between these attention processes. That is, interference between endogenous and exogenous attention occurs whenever both attention systems recruit the same sensory processes. These limitations in processing likely follow from conflicting patterns of neuronal excitation and inhibition whenever both engage opposite sides of the visual field. Indeed, whenever endogenous attention alters the excitability of neuronal populations involved in sensory processing at the contralateral side, an effect captured by alpha-band activity, this process is disrupted by exogenous attention producing neuronal inhibition in the same neuronal population. Our findings therefore support the notion that exogenous and endogenous attention recruit the same neural resources for visual processing and that the limited processing resources of the brain that govern the dynamics between exogenous and endogenous attention.

## Methods

### Participants

We recruited 42 participants based on convenience sampling. Participants received monetary compensation of $10/hour for completing 4 different tasks. Each task consisted of 384 trials. Participants performed 10 practice trials before each task. We analyzed data from 3 tasks: the non-cueing task, the endogenous cueing task, and the double cueing task. The experiment was a single testing session and lasted approximately 2 hours. All participants had normal or corrected-to-normal vision and provided consent. The experimental protocol was approved by the McGill Ethics Board Office.

Seven participants were excluded due to poor EEG data quality. One participant did not complete all tasks. We additionally excluded one participant due to below chance discrimination accuracy rate (~40% averaged across all conditions) and one participant due to a high volume of timeout errors (response times > 1500ms on ~11% of trials overall). The final sample included 32 individuals (24 women, Mean age = 21.9 (SD = 2.6)). Our sample size follows from previous work using the double cueing experimental approach: Effect size estimates of task performance suggest that 6 participants are required to achieve a power of .8 for the within-individual cueing effects of exogenous and endogenous orienting in a target discrimination task for alpha = .05 (Landry, Da Silva Castanheira, Sackur, & Raz, 2021). The sample size of the current study is therefore adequate to detect the effect of attention at the behavioral level. Our sample size is almost twice that of previous EEG experiments investigating exogenous and endogenous orienting together (Hopfinger & West, 2006; Keefe & Störmer, 2021).

### Stimuli, Apparatus & Design

Participants viewed stimuli on a 24-in BenQ G2420HD monitor sitting approximately 60 cm away. Stimulus presentation was done using MATLAB R2015b (Mathworks Inc., Natick, MA, USA) and the third version of the Psychophysics toolbox (Brainard & Vision, 1997; Kleiner et al., 2007; Pelli, 1997). The screen was set to 75hz. Except for the target, all stimuli were black (i.e., RGB values of 0, 0, 0; 1.11 cd/m2) and white (i.e., RGB values of 255, 255, 255; 218.8 cd/m2) drawings on a grey background (i.e., RGB values of 128, 128, 128; 70.88 cd/m2). The fixation marker was an empty circle made from a black line drawing with a radius of 1.2° located in the center of the screen. Two target placeholders were located at 8.7° on each side of the fixation marker on the left and right side of the screen. These placeholders were made from black line drawings of empty circles with a 2.4° radius. We cued exogenous attention by briefly changing the line drawing from one of the placeholders to white. To ensure that this cue solely engaged exogenous orienting, the cue-target spatial contingency was set to 50%, such that the cue was only predictive of the target’s location at chance-level. The exogenous cue was therefore non-informative and occurred at the periphery. This is consistent with the standard approach for eliciting exogenous attention in the lab (Chica et al., 2014). We cued endogenous attention by coloring the inside of the fixation marker, wherein the right or left half of the circle was shaded in black, and the other half in white. The side of the fixation marker that turned white indicated where the target was likely to occur. For example, if the right side turned white, the target was 66.6% likely to appear in the right placeholder. In this way, we avoided using overlearned directional cues, like an arrow, to engage endogenous attention (Ristic & Landry, 2015). Participants were aware of these contingencies. The targets were sinusoidal black and white gratings combined with a Gaussian envelope. Spatial frequency was set to 3 cpd. Target stimuli were tilted 5° degrees clockwise or counter-clockwise.

### Procedure

Participants completed four tasks of 384 trials: A non-cueing task, an exogenous cueing task, an endogenous cueing task, and a double cueing task where both exogenous and endogenous orienting were engaged. Task order was randomized across participants. Note that the current manuscript includes data from the non-cueing, the endogenous cueing, and the double cueing tasks.

Participants stared at the center of the screen throughout the experiment, while we assessed their eye movements using electro-oculogram. We jittered the latencies between fixation and attention cues, between both attention cues, and between the cues and the target. We used a uniform distribution of latencies to minimize the effects of temporal prediction following spatial cueing. Cue-target latencies were adjusted following the temporal profiles of exogenous and endogenous attention so that the benefit of attention processing would be optimized for both attention (Chica et al., 2014).

In the non-cueing task, the timing between the fixation circle and target stimulus was jittered from 1027ms to 1280ms. The target stimulus stayed on the screen until participants responded. We used the non-cueing task as a baseline against which we regressed the sensory effects of cue stimuli for the target-related analysis (see the *Electroencephalography* section below).

In the endogenous orienting task, again the timing between the fixation circle and target stimulus was jittered between 1027ms and 1280ms, followed by the endogenous cue that remained on the screen until participants responded. The timing between the onset of the endogenous cue and target onset was jittered between 613ms and 926ms. The target stimulus remained on screen until participants responded (Figure 1).

The double cueing task was similar to the single cueing task, with the exception that we interleaved a peripheral cue between the endogenous cue and the target events. The timing between the fixation circle and the target event was again jittered between 1027ms and 1280ms. The timing between the endogenous cue and the target stimulus varied between 613ms and 926ms. Next, the peripheral cue would onset and then quickly offset after 106ms. The timing between the onset of peripheral cue and the target stimulus was jittered between 106ms and 307ms.

Participants were instructed to complete a target discrimination task and indicate the orientation of the Gabor target as quickly and accurately as possible on a QWERTY keyboard by pressing the F key for counter-clockwise orientation and the J key for clockwise orientation. Inter-trial period was set to 1s.

### Data analyses for behavioral performance

Participants’ discrimination performance was near ceiling; the average accuracy rate was ~93%. Thus, we examined task performance via accurate response times (RTs). We removed trials where participants made anticipation (i.e., RTs < 150ms) or timeout (i.e., RTs > 1500ms) errors. This accounted for less than 2% of trials overall. We additionally removed wrong key presses, which corresponded to less than 1% of trials. We used hierarchical linear regression models (Gelman & Hill, 2006) to test the effects of cueing (i.e., valid versus invalid) and task (i.e., single versus double cueing). Fixed factors were added in stepwise fashion while we used a chi-square goodness-of-fit test over the deviance to determine whether they significantly improved the fit. We computed the Bayesian information criterion (BIC) to select the most parsimonious model. Lastly, we calculated Bayes factors to weight evidence for the alternative against the null hypothesis based on the BIC approximation (Wagenmakers, 2007) :

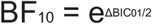

For the single cueing endogenous task, we included cue validity as a fixed factor. For the double cueing task, we included endogenous cue validity, exogenous cue validity, and their interaction as fixed factors. Given that our study aimed to compare single and double cueing conditions, we examined if the cueing effect of endogenous attention varied as a function of cueing condition (i.e., single and double cueing), wherein we included cue validity (i.e., endogenous cue valid versus endogenous cue invalid) and task (i.e., single versus double cueing) as fixed factors. For all regression models, we added participants as a random factor to account for the within-subject design of the study.

### Electroencephalography

We recorded EEG signals using 64 Ag/AgCl active electrodes at a sampling rate of 1000 Hz (ActiCap System; Brain Products GmbH; Gilching, Germany). We monitored eye blinks and eye movements through additional bipolar electrodes placed at the outer canthi, as well as the superior and inferior orbits of the left eye. We kept impedances of all electrodes below 10 kΩ, while all electrophysiological signals were amplified (ActiChamp System; Brain Products GmbH; Gilching, Germany). Electrodes were referenced online to C4. We re-referenced the electrodes offline to the average of all channels. Preprocessing and analysis were conducted in BrainVision Analyzer (ActiChamp System; Brain Products GmbH Inc.; Gilching, Germany) and MATLAB (R2020a; Mathworks Inc., Natick, MA) using Brainstorm (Tadel, Baillet, Mosher, Pantazis, & Leahy, 2011) and custom scripts. We downsampled the data to 250 Hz and visually inspected EEG signals to identify activity exceeding ± 200 μV. We applied two IIR Butterworth filters: a first High-pass 0.1 Hz filter of 8th order and a 60Hz notch filter. We interpolated bad channels topographically (1.12% of channels) and then identified artifacts related to eye movements and blinks using independent component analysis via the BrainVision Analyzer Ocular correction ICA tool.

For target-related events, we removed event-related responses from exogenous and endogenous cues by regressing-out the effects of the task. Here we took the residuals of the linear regression model using the non-cueing condition as baseline. A hierarchical linear regression was run in MATLAB with cueing condition as a dummy coded fixed factor, and subjects as a random factor. ‘Raw’ Residuals were obtained from all channels and subjects.

### Event-related potentials

We analyzed target-related event-related potentials (ERPs). We first applied a FIR bandpass-pass filter between 0.5 and 15 Hz and then divided the EEG into epochs spanning −200 and 1000ms. All triggers were realigned according to a photodiode stimulation linked to the onset of the target event. ERPs were baseline corrected from −100 to 0ms.

### Lateralized alpha index

We computed the lateralized alpha index (LAI) based on time-resolved alpha power. These neural rhythms were computed for target-locked epochs for every channel from −1000ms and 500ms. We first estimated the instantaneous amplitude of alpha waves (i.e., 8 to 12hz) based on the real part of the complex filter Hilbert transformation. Next, we computed the LAI following the location of the direction of the endogenous attention at each time point based on to the following equation:

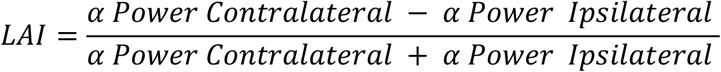

In one series of analysis, we defined contra- and ipsilateral components relative to the direction of endogenous orienting. In another, we defined it relative to the location of the target event. See multivariate analyses and hierarchical linear regression modelling sections for details. We computed LAI for lateral channels only and excluded central ones. Computations were done such that every lateral channel was paired with its corresponding match on the ipsilateral side. Since we were interested in pre-stimulus alpha power, we baseline corrected the LAI from 200ms to 300ms post-target stimulus.

### Multivariate analyses

We leveraged multivariate statistical techniques to evaluate the influence of exogenous attention on the cueing effects of endogenous attention. Our approach was twofold. In our first approach, we trained classifiers to decode the effects of endogenous cue validity during single cueing and then evaluated their performance in the context of double cueing (Figure 2A). Therefore, this first approach evaluated the classifiers’ capacity to decode the cueing effects of endogenous attention alone based on target-locked ERPs. We then assessed whether the performance of these classifiers is impaired during double cueing. We reasoned that classification should hardly vary across cueing conditions if exogenous attention hardly impacts the endogenous cueing effect for target processing. Conversely, a significant drop in classification performance would indicate that exogenous attention alters the cueing effect of endogenous attention. In our second approach, we trained and tested classifiers in their ability to decode cueing conditions -- i.e., single versus double cueing conditions (Figure 2B). Here, we aimed to see whether endogenous attention varied from single to double cueing. In this fashion, we trained and tested SVM classifiers in their ability to accurately separate single vs. double cueing congruent and single vs. double cueing incongruent.

Our multivariate approach largely follows from the work of Bae and Luck (2018). Here, we performed multivariate classification using linear support vector machine (SVM) and MATLAB’s *fitcsvm* and *predict* functions. The training and testing phases were completed at the participant’s level. Our first analysis aimed to assess how classifiers trained to decode the cue validity effect of endogenous attention (i.e., endogenous cue valid vs. endogenous cue invalid) along the time series of target-locked ERP from the single cueing condition would perform when they are tested in the context of double cueing. Here, we tested the performance of the SVM models separately for congruent and incongruent cueing trials. We used a three-fold cross-validation procedure wherein trials from the single cueing condition were separated into three different bins per class – i.e., three bins of trials for endogenous cue valid and three for endogenous cue invalid. We equated trials per bins for all conditions and participants. There were 14 trials per bin. We applied multivariate classification to left and right target location separately because we did not want the target’s location to be a confounding feature. Target location was therefore orthogonal to the classifiers’ performance. We decoded the cueing effect of endogenous attention from target-locked ERP by averaging the EEG from each bin, and then baseline corrected them from −100ms to 0ms. Then, we trained SVM on each time point along the time series by including the EEG channels as features in our model. We trained the classifier using two ERP from each class (i.e., two ERP for invalid cueing and two for valid cueing), and then tested the model on the remaining ones (i.e., one ERP for invalid cueing and one for valid cueing). We iterated this process such that each ERP was used twice for training and once for testing. We repeated this process 50 times per participant while randomly shuffling trials across bins for each repetition. Lastly, we averaged classification accuracy rates for each participant across both target locations and smoothed these values across the time series via a five-sample sliding window -- i.e., a 20ms window (Hong et al., 2020). This procedure allowed us to assess the classification performance across the time series during single cueing.

Our first analysis therefore assessed the performance of SVM classifiers that were trained to decode endogenous cue validity during single cueing. Here, we tested these classifiers’ ability to classify endogenous cue validity in three separate contexts: (1) on the left-out data from single cueing condition; (2) on the data from the double cueing condition for congruent cueing trials; and (3) on the data from the double cueing condition for incongruent cueing trials. We used one sample t-tests across the time series to determine whether classification performances were better than chance-level. We also compared classification accuracies using pairwise t-tests across the time series: A first one where we contrasted the decoding accuracy of single cueing against that of double cueing congruent, a second one where we contrasted the decoding accuracy from single cueing against that of double cueing incongruent, and lastly, we compared decoding accuracies from double cueing congruent and incongruent. For all analyses, we controlled for family-wise errors via random permutations and mass t-statistics for cluster size. The cluster forming threshold was set to p < .05. We performed 1000 permutations where we randomly varied the classification labels and then contrasted observed cluster sizes based on t-statistics against surrogate distributions of cluster size. Statistically significant clusters were threshold at ≥ 95%.

For our second analysis, we examined whether SVM classification could decode the cueing condition (i.e., single vs. double cueing) based on interference patterns from exogenous attention over the cueing effect of endogenous attention by training SVM classifiers to decode cueing conditions (i.e., single versus double cueing) based on target-locked waveforms for the cueing effect of endogenous attention (i.e., endogenous valid minus endogenous cue invalid). Again, we trained and tested the classifiers across the time series while channels were included as features in our model. Note that we performed separate analyses for contralateral and ipsilateral channels. We relied on the same three-fold cross-validation procedure we previously described, while we grouped trials into separate bins for valid and invalid trials across single and double cueing. Each bin comprised 14 trials. We then averaged the trials from each bin and then subtracted the waveform of the first invalid bin from that of the first valid bin separately for single and double cueing, likewise for the second and third invalid and valid bins. This process yielded three waveforms for single cueing and three for double cueing, with each waveform corresponding to the cueing effect of endogenous attention (i.e., endogenous cue valid minus endogenous cue invalid). Again, our training and validation procedures were identical to the ones we described previously (Bae & Luck, 2018), wherein we trained the SVM classifier on 2 waveforms from each cueing condition (i.e., single versus double cueing) and validated the remaining one. Once more, we iterated this procedure such that each waveform was used twice for training and once for testing and repeated this process 50 times for each participant, while randomly selecting the trials for each bin on every repetition. Overall, we applied this multivariate classification procedure four times: When both cues indicated the same location and then again when they indicated opposite locations, both across contra- and ipsilateral components. For all procedures, we verified whether decoding accuracy rates across participants was greater than chance-level (i.e., 50%) using cluster-corrected one-sample t-tests. Lastly, to better understand the encoding model, we used the method described by Haufe et al. (2014) and multiplied the weights of the classifiers with the covariance matrix of the neural data along the time series.

We used the same approach where we solely examined endogenous processing of target events. Instead of examining the waveform related to the endogenous cueing effect (i.e., endogenous cue valid minus endogenous cue invalid), we solely focused on trials where the endogenous cue was valid. In this way, we were able to test whether the concurrent engagement of exogenous attention impacts processing of the target event following orienting of endogenous attention.

We adopted a similar approach to examine the modulations of LAI across single and double cueing using multivariate statistics. Here, we computed target-locked LAI during the pre-stimulus period following the direction of endogenous orienting. LAI time series were baseline corrected from 200ms and 300ms post-stimulus. Again, we separated analyses from the double cueing condition when both cues pointed towards the same location and when both cues indicated opposite locations. For this analysis, we separated trials of instantaneous alpha power into bins of contralateral and ipsilateral channels. Each bin comprised 32 trials. Next, we computed the averaged from each bin and computed the LAI by matching the contra- and ipsilateral components of the first, second and third bins across single and double cueing conditions. Lastly, we adopted the same three-fold cross-validation as before to evaluate the effects of the cueing condition over LAI.

### ERPs of visual processing

We extracted target-locked ERPs at channel PO7 and PO8 based on previous work showing that visuospatial attention modulates early visual components of the target-locked waveforms. Here, we extracted the waveform at the contralateral channel relative to the target location for each trial. For each participant, we averaged the waveforms at the contralateral channel separately for the following experimental conditions: Single cueing endogenous valid trials; single cueing endogenous invalid trials; double cueing opposite endogenous valid trials; double cueing opposite endogenous invalid trials. Next, we tested whether the onset of the exogenous to the opposite location relative to that of the endogenous cue affect endogenous cue validity effect for early EEG waveforms using hierarchical regression models at each time point where we included endogenous cue validity (endogenous cue valid vs. endogenous cue invalid), cueing condition (single vs. double cueing), as well as their interaction as fixed factor, wihle participants were included as random factors:

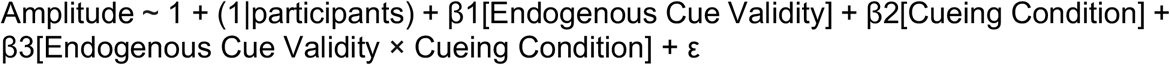

We cluster-corrected across time using mass t-statistics.

### Moderated Mediation Analysis

To test our hypothesis that exogenous attention interferes with endogenous attention through alpha rhythms, we first identified the channel and the time where the SVM beta weight was maximal to discriminate between single and double cueing based on LAI when both cues indicate opposite locations during double cueing. We observed the maximal SVM weight value for channel P3/ P4 at 40ms post-target onset. We therefore extracted the LAI value at that specific channel and time point to evaluate our hypothesis. We relied on a moderated mediation analysis to assess our hypothesis, wherein changes in LAI mediates the relationship between endogenous cue validity and RTs, and the onset of the exogenous cue moderates this mediation effect. We validated our assumption that LAI mediates the relationship between endogenous cue validity and RTs before testing our moderation effect. Here, we used a simple mediation model based on the following regression models:

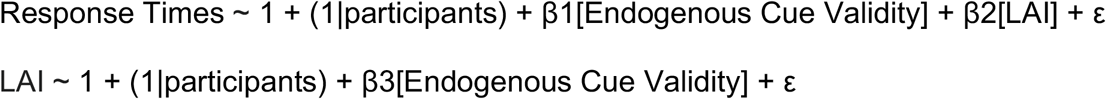

We used the Sobel approach to estimate the indirect effect and computed the product of regression coefficients β2*β3 (Sobel, 1982). We computed the p-value by comparing the indirect effect coefficient to a surrogate null distribution, which we obtained via 1000 permutations where we randomly shuffled the LAI values and endogenous cue validity dummy variable. The p-value for the indirect was smaller than .001, which validated our assumption (supplementary Figure 4; supplementary Tables 7 and 8). Next, we evaluated our main hypothesis and tested whether the cueing condition (single versus double cueing) moderates this mediation effect. Our previous analysis (see Behavior section) highlighted a significant interaction between endogenous cue validity and cueing condition (single versus double cueing) for predicting RTs when both cues indicated opposite locations during double cueing. This interaction reflects the interference effect of exogenous attention over endogenous cue validity. We therefore aimed to see if the moderation of the mediation effect via LAI explains this interference effect. Thus, rather than aiming to explain the direct relationship between endogenous cue validity and RTs, we aimed to explain the interaction between endogenous cue validity and cueing condition. Again, we adopted the Sobel approach based on the following regression models (see supplementary Tables 11 and 12):

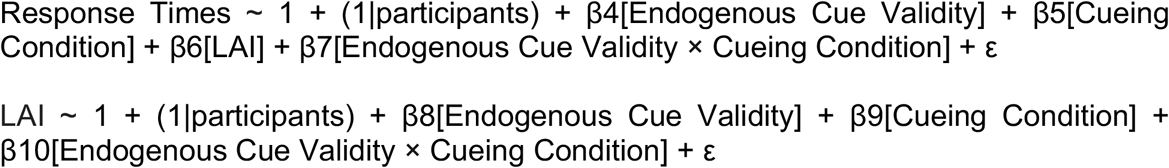

We estimated the moderated mediation index based on the product of coefficients β4*β10+β6*β10 (Hayes, 2015; Preacher, Rucker, & Hayes, 2007). We tested significance based on 10,000 bootstraps with replacement of the parameters coefficients.

## Acknowledgments

M.L. acknowledges support from the Fonds de Recherche du Québec – Nature et Technologies. J.D.C. acknowledges a doctoral fellowship from the Natural Science and Engineering Research Council of Canada. This work was supported by a grant from CIHR and tiers-2 Canada Research Chair to A.R. S.B. is supported by the NIH (R01 EB026299), a Discovery grant from the Natural Sciences and Engineering Research Council of Canada (436355-13), the CIHR Canada research Chair in Neural Dynamics of Brain Systems, the Brain Canada Foundation with support from Health Canada, and the Innovative Ideas program from the Canada First Research Excellence Fund, awarded to McGill University for the Healthy Brains for Healthy Lives initiative (1c-II-14). J.S. acknowledges grants from the ANR (France): ANR-16-ASTR-0014 and ANR-17-EURE-0017. We would like to thank Anna Prokusheva for her help collecting the data. Lastly, we would also like to thank Barry Giesbrecht, Mathieu Roy, Dave Saint-Amour, Vincent de Gardelle, Viola Störmer for helpful comments about this research project.

