## Supplementary Material for "Exogenous attention interferes with endogenous attention processing via lateralized alpha power"

We present the tables from all hierarchical linear regression models for predicting correct response times. We performed the comparison and selection of models using chi-square goodness-of-fit tests and Bayesian Information Criterion (BIC). The best fitting model is highlighted in bold. Model comparison was done in a stepwise manner.

**Table 1.** Single endogenous cueing task. Hierarchical linear regression models for predicting RTs as a function of endogenous cue validity.

| Models | $X^2$ | p-value | BIC |
|---|---|---|---|
| RTs ~ β0 + S0 [*Subject*] + ε | | | 147240 |
| **RTs ~ β0 + S0[*Subject*] + β1[*Endo.CueValidity*] + ε** | **126.04** | **p < .001** | **147130** |

**Table 2.** Single endogenous cueing task. Hierarchical linear regression parameters of the best model for predicting RTs as a function exogenous cue validity.

Model: RTs ~ 1 + (1|participant) + β1[*Endo.CueValidity*] + ε

| Variables | Coefficient | Std. Error | 95% C.I. |
|---|---|---|---|
| *Intercept* | 649.53 | 14.95 | [620.23, 678.84] |
| *Endo Cue Validity* | -36.49 | 3.24 | [-42.84, -30.13] |

**Table 3.** Double cueing task. Hierarchical linear regression models for predicting RTs as a function of exogenous and endogenous cue validity.

| Models | $X^2$ | p-value | BIC |
|---|---|---|---|
| RTs ~ 1 + (1\|participant) + ε | | | 145300 |
| RTs ~ 1 + (1\|participant) + β1[*Exo.CueValidity*] + ε | 166.27 | p < .001 | 145150 |
| **RTs ~ 1 + (1\|participant) + β1[*Exo.CueValidity*] + β2[*Endo.CueValidity*] + ε** | **48.37** | **p < .001** | **145110** |
| RTs ~ 1 + (1\|participant) + β1[*Exo.CueValidity*] + β2[*Endo.CueValidity*] + β3[*Exo.CueValidity* × *Endo.CueValidity*] + ε | < .1 | p = .98 | 145120 |

**Table 4.** Single cueing task. Hierarchical linear regression parameters of the best fitting model for predicting RTs as a function exogenous and endogenous cue validity.

Model: RTs ~ 1 + (1|participant) + β1[*Exo.CueValidity*] + β2[*Endo.CueValidity*] + ε

| Variables | Coefficient | Std. Error | 95% C.I. |
|---|---|---|---|
| *Intercept* | 670.72 | 16.36 | [638.66, 702.78] |
| *Exo Cue Validity* | -39.27 | 3.04 | [-45.22, -33.32] |
| *Endo Cue Validity* | -22.41 | 3.21 | [-28.73, -16.11] |

**Table 5.** Comparing single to double cueing – same cueing location. Hierarchical linear regression models for predicting RTs as a function of endogenous cue validity and cueing condition.

| Models | $X^2$ | p-value | BIC |
|---|---|---|---|
| RTs ~ 1 + (1\|participant) + ε | | | 220860 |
| RTs ~ 1 + (1\|participant) + β1[*Endo.CueValidity*] + ε | 274.84 | p < .001 | 220590 |
| RTs ~ 1 + (1\|participant) + β1[*Enddo.CueValidity*] + β2[*Cueing condition*] + ε | 2.94 | p = .09 | 220600 |
| **RTs ~ 1 + (1\|participant) + β1[*Endo.CueValidity*] + β2[*Task*] + β3[*Endo.CueValidity × Cueing condition*] + ε** | **19.67** | **p < .01** | 220590 |

We additional compared the 2nd and 4th models, which indicated a significant difference $X^2$=22.60, p < .001.

**Table 6.** Comparing single to double cueing – same cueing location. Hierarchical linear regression parameters of the best models for predicting RTs as a function of exogenous cue validity and cueing condition.

Model: RTs ~ 1 + (1|participant) + β1[*Endo.CueValidity*] + β2[*Cueing Condition*] + ε

| Variables | Coefficient | Std. Error | 95% C.I. |
|---|---|---|---|
| *Intercept* | 649.1 | 14.24 | [621.19, 677.02] |
| *Endo Cue Validity* | -36.2 | 3.27 | [-42.61, -29.79] |
| *Cueing condition* | 21.42 | 4.64 | [12.32, 30.52] |

| | | | |
|---|---|---|---|
| *Endo Cue Val. × Cueing condition* | -25.2 | 5.68 | [-36.33, -14.1] |

**Table 7.** Comparing single to double cueing – opposite cueing locations. Hierarchical linear regression models for predicting RTs as a function of endogenous cue validity and cueing condition.

| Models | $X^2$ | p-value | BIC |
|---|---|---|---|
| RTs ~ 1 + (1\|participant) + ε | | | 219890 |
| RTs ~ 1 + (1\|participant) + β1[*Endo.CueValidity*] + ε | 48.92 | p < .001 | 219850 |
| RTs ~ 1 + (1\|participant) + β1[*Enddo.CueValidity*] + β2[*Cueing condition*] + ε | 41.79 | p < .001 | 219820 |
| **RTs ~ 1 + (1\|participant) + β1[*Endo.CueValidity*] + β2[*Task*] + β3[*Endo.CueValidity × Cueing condition*] + ε** | **85.55** | **p < .001** | 219740 |

**Table 8.** Comparing single to double cueing – opposite cueing locations. Hierarchical linear regression parameters of the best models for predicting RTs as a function of exogenous cue validity and cueing condition.

Model: RTs ~ 1 + (1|participant) + β1[*Endo.CueValidity*] + β2[*Cueing Condition*] + ε

| Variables | Coefficient | Std. Error | 95% C.I. |
|---|---|---|---|
| *Intercept* | 649.29 | 14.65 | [620.58, 677.99] |
| *Endo Cue Validity* | -36.28 | 3.29 | [-42.73, -29.83] |
| *Cueing condition* | -17.55 | 4.67 | [-26.71, -8.39] |
| *Endo Cue Val. × Cueing condition* | 52.74 | 5.72 | [41.52, 63.97] |

We used multivariate pattern classification to examine endogenous cue validity across single and double cueing. Here, we show that the classifiers were able to perform above chance level (i.e., 50%) for single cueing and for double cueing when both cues indicated the same location. In turn, however, we observed that classification was below chance-level in the double cueing condition whenever both cues indicated opposite locations.

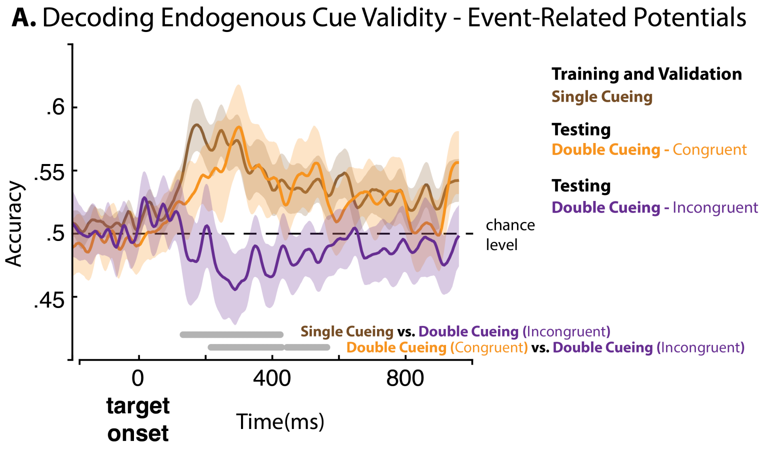


**Supplementary Figure 1.** Multivariate classification of endogenous cue validity based on cue-locked ERPs where we trained a SVM linear classifier in the context of single cueing and tested it in contexts of single (**A**) and double cueing (**B**, **C**). We separated double cueing into trials where both cues were indicating the same location and trials where they indicated opposite locations. Classification was done at subject-level. We compared classification performances across all condition using cluster-corrected pairwise t-tests against chance-level (i.e., 50%). Grey lines indicate significant clusters. Error bars indicate the SEM for each condition.

We further explored the relationship between cueing condition, endogenous cue validity, LAI and response times. In this effort, we used a moderated mediating analysis. However, we first tested a simple mediation. Supplementary tables 9 and 10 show regression coefficients for this analysis. Next, we tested the moderated mediation effect. Supplementary tables 11 and 12 show regression coefficients for this analysis.

**Table 9.** Hierarchical linear regression parameters for predicting RTs as a function of endogenous cue validity and LAI

Model: Response Time ~ 1 + (1|participant) + β1[*Endo Cue Validity*] + β2[*LAI*] + ε

| Variables | Coefficient | Std. Error | 95% C.I. |
|---|---|---|---|
| *Intercept* | 651.83 | 15.74 | [620.69, 682.97] |
| *Endo Cue Validity* | -26.7 | 8.94 | [-44.4, -9.02] |
| *LAI* | 114.56 | 36.23 | [44.86, 186.27] |

**Table 10.** Hierarchical linear regression parameters for predicting RTs as a function of endogenous cue validity

Model: LAI ~ 1 + (1|participant) + β3[*Endo Cue Validity*] + ε

| Variables | Coefficient | Std. Error | 95% C.I. |
|---|---|---|---|
| *Intercept* | -.07 | .01 | [-.1, -.04] |
| *Endo Cue Validity* | .10 | .02 | [.06, .15] |

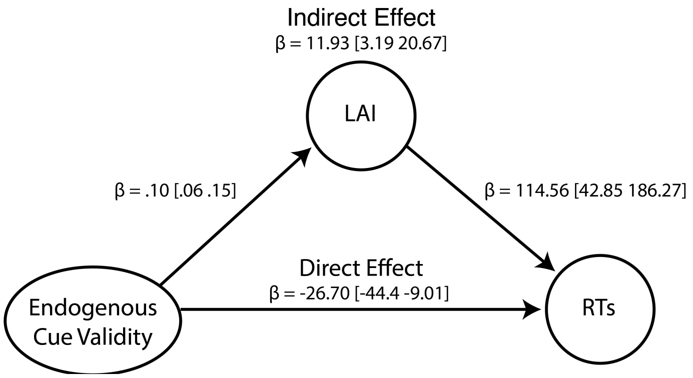


**Supplementary Figure 4.** Mediation analysis where the direct effect involves endogenous cue validity predicted response times, wherein LAI mediates this relationship.

**Table 11.** Hierarchical linear regression parameters for predicting RTs as a function of endogenous cue validity

Model: Response Times ~ 1 + (1|participant) + β4[*Endo Cue Validity*] + β5[*Cueing Condition*] + β6[LAI] + β7[*Endo Cue Validity × Cueing Condition*] + ε

| Variables | Coefficient | Std. Error | 95% C.I. |
|---|---|---|---|
| *Intercept* | 652.64 | 16.61 | [619.78, 685.51] |
| *Endo Cue Validity* | -34.51 | 12.65 | [-59.54, -9.48] |
| *Cueing Condition* | -4.05 | 13.13 | [-30.03, 21.93] |
| *LAI* | 97.03 | 43.85 | [10.23, 183.83] |
| *Endo Cue Validity × Cueing Condition* | 16.21 | 20.21 | [-23.95, 56.38] |

**Table 12.** Hierarchical linear regression parameters for predicting RTs as a function of endogenous cue validity

Model: LAI ~ 1 + (1|participant) + β8[*Endo Cue Validity*] + β9[*Cueing Condition*] + β10[*Endo Cue Validity × Cueing Condition*] + ε

| Variables | Coefficient | Std. Error | 95% C.I. |
|---|---|---|---|

| | | | |
|---|---|---|---|
| *Intercept* | .01 | .02 | [-.03, .04] |
| *Endo Cue Validity* | -.03 | .03 | [-.08, .03] |
| *Cueing Condition* | -.15 | .03 | [-.2, -.1] |
| *Endo Cue Validity × Cueing Condition* | .24 | .04 | [.17, .31] |
